# Timescale and genetic linkage explain the variable impact of defense systems on horizontal gene transfer

**DOI:** 10.1101/2024.02.29.582795

**Authors:** Yang Liu, João Botelho, Jaime Iranzo

**Affiliations:** Centro de Biotecnología y Genómica de Plantas, Universidad Politécnica de Madrid (UPM) - Instituto Nacional de Investigación y Tecnología Agraria y Alimentaria (INIA-CSIC), Madrid, Spain; Centro de Astrobiología (CAB), CSIC-INTA, Madrid, Spain; Institute for Biocomputation and Physics of Complex Systems (BIFI), University of Zaragoza, Zaragoza, Spain

**Keywords:** defense system, horizontal gene transfer, mobile genetic element, CRISPR-Cas, phage, plasmid, comparative genomics

## Abstract

Prokaryotes have evolved a wide repertoire of defense systems to prevent invasion by mobile genetic elements (MGE). However, because MGE are vehicles for the exchange of beneficial accessory genes, defense systems could consequently impede rapid adaptation in microbial populations. Here, we study how defense systems impact horizontal gene transfer (HGT) in the short and long terms. By combining comparative genomics and phylogeny-aware statistical methods, we quantified the association between the presence of 7 widespread defense systems and the abundance of MGE in the genomes of 196 bacterial and 1 archaeal species. We also calculated the differences in the rates of gene gain and loss between lineages that possess and lack each defense system. Our results show that the impact of defense systems on HGT is highly taxon- and system-dependent. CRISPR-Cas stands out as the defense system that most often associates with a decrease in the number of MGE and reduced gene acquisition. Timescale analysis reveals that defense systems must persist in a lineage for a relatively long time to exert an appreciable negative impact on HGT. In contrast, at short evolutionary times, defense systems, MGE, and gene gain rates tend to be positively correlated. Based on these results and given the high turnover rates experienced by defense systems, we propose that the inhibitory effect of most defense systems on HGT is masked by recent co-transfer events involving MGE.

## Introduction

Gene exchange plays a key role in the adaption of microbes to changing environments, facilitating the spread of antibiotic resistance, pathogenicity factors, metabolic genes, and other accessory functions (Arnold et al. 2022). Over the last decade, there has been an increasing interest in assessing the ecological and genetic factors that control horizontal gene transfer (HGT) and determine the outcome of newly acquired genes in microbial populations (Soucy et al. 2015; Hall et al. 2020; Lee et al. 2022). HGT is often mediated by mobile genetic elements (MGE), such as phages, integrative and conjugative elements, and plasmids, against which bacteria have evolved an elaborate repertoire of defense systems (Doron et al. 2018; Botelho et al. 2023; Georjon and Bernheim 2023; Mayo-Munoz et al. 2023; Shaw et al. 2023). As a result, large-scale patterns of HGT are shaped by an interplay of ecological and genetic variables that underlie cross-strain and cross-species differences in susceptibility to MGE (Haudiquet et al. 2022).

Recent studies have highlighted the role of CRISPR-Cas as widespread adaptive immunity systems that protect archaea and bacteria against viruses and other MGE (van Vliet et al. 2021; Watson et al. 2024). While in vitro experiments have provided supportive evidence for this function (Marraffini and Sontheimer 2008; O’Hara et al. 2017; Watson et al. 2018), questions remain about the extent to which CRISPR-Cas systems constrain gene exchange in nature (Gophna et al. 2015; O’Meara and Nunney 2019; Shehreen et al. 2019; Westra and Levin 2020; Wheatley and MacLean 2021; Pursey et al. 2022). More generally, there is a paucity of research on how other defense systems, such as restriction-modification (RM), abortive infection (Abi), and an expansive repertoire of recently identified gene systems including Gabija, CBASS (cyclic oligonucleotide-based antiphage signaling system), DMS (DNA modification-based systems), and DRT (defense-associated reverse transcriptases) affect HGT in prokaryotes (Tesson et al. 2022; Costa et al. 2024).

In contrast with the expectation that defense systems restrict HGT by interfering with the propagation of MGE, several empirical and theoretical observations suggest that the relation between defense systems and HGT might be more complex. First, the selection pressure to maintain defense systems in a population (and consequently their prevalence) generally increases with the exposure to MGE (Oliveira et al. 2016; Meaden et al. 2022). Second, defense systems are mobilized by MGE, which could lead to a trivial positive association with HGT rates. In a less trivial manner, whole defense systems or parts of them are often encoded by MGE (Makarova et al. 2011; Pinilla-Redondo et al. 2022; Botelho 2023), promoting the retention of the latter for their beneficial side-effects in cellular defense (Koonin et al. 2020; Rocha and Bikard 2022).

Here, we investigated the association between 7 widespread defense systems and HGT rates in 197 prokaryotic species. By combining high-quality genomic data, phylogenomic methods, and phylogeny-aware statistical inference, our study aimed to shed light on the nuanced consequences of the interplay between defense systems and MGE on bacterial evolution across time scales.

## Results

### Association between defense systems and MGE abundance is MGE- and taxon-dependent

We used species-wise phylogenetic generalized linear mixed models (PGLMM) to study the association between the presence or absence of the 7 most prevalent defense systems in the dataset (RM, DMS, Abi, CRISPR-Cas, Gabija, DRT, and CBASS), genome size (measured as the total number of genes), and the number of MGE per genome (Fig. 1, Tables S1-S4). Notably, the sign and magnitude of the associations are strongly taxon-, system-, and MGE-dependent. Out of 197 species included in the analysis, around 20-30 (depending on the defense system) displayed a statistically significant (*p* < 0.05) positive association between the presence of the defense system and genome size. In contrast, statistically significant negative associations were only observed in 5-15 species (Fig. 1a, top row). Because differences in the number of genomes per species could bias comparisons based on p-values, we also explored an alternative criterion based on effect sizes to determine the number of positive and negative associations (see Methods). Regardless of the criterion, CRISPR-Cas stood out as the defense system that most often displays negative associations with genome size. In contrast, other defense systems positively correlate with larger genomes in most of the species. The analysis also reveals differences regarding the association between defense systems and distinct types of MGE (Fig. 1a, middle rows). In all defense systems, negative associations with prophages are more frequent than negative associations with plasmids. Once again, the clearest trend corresponds to CRISPR-Cas, whose presence often correlates with a reduction in the number of prophages, but with higher numbers of transposable elements and plasmids. Significant associations, when detected, affect a sizeable fraction of the accessory genome, with 20-40% differences in MGE content between genomes that do and do not harbor the defense system (Fig. 1b).

**Figure 1:**
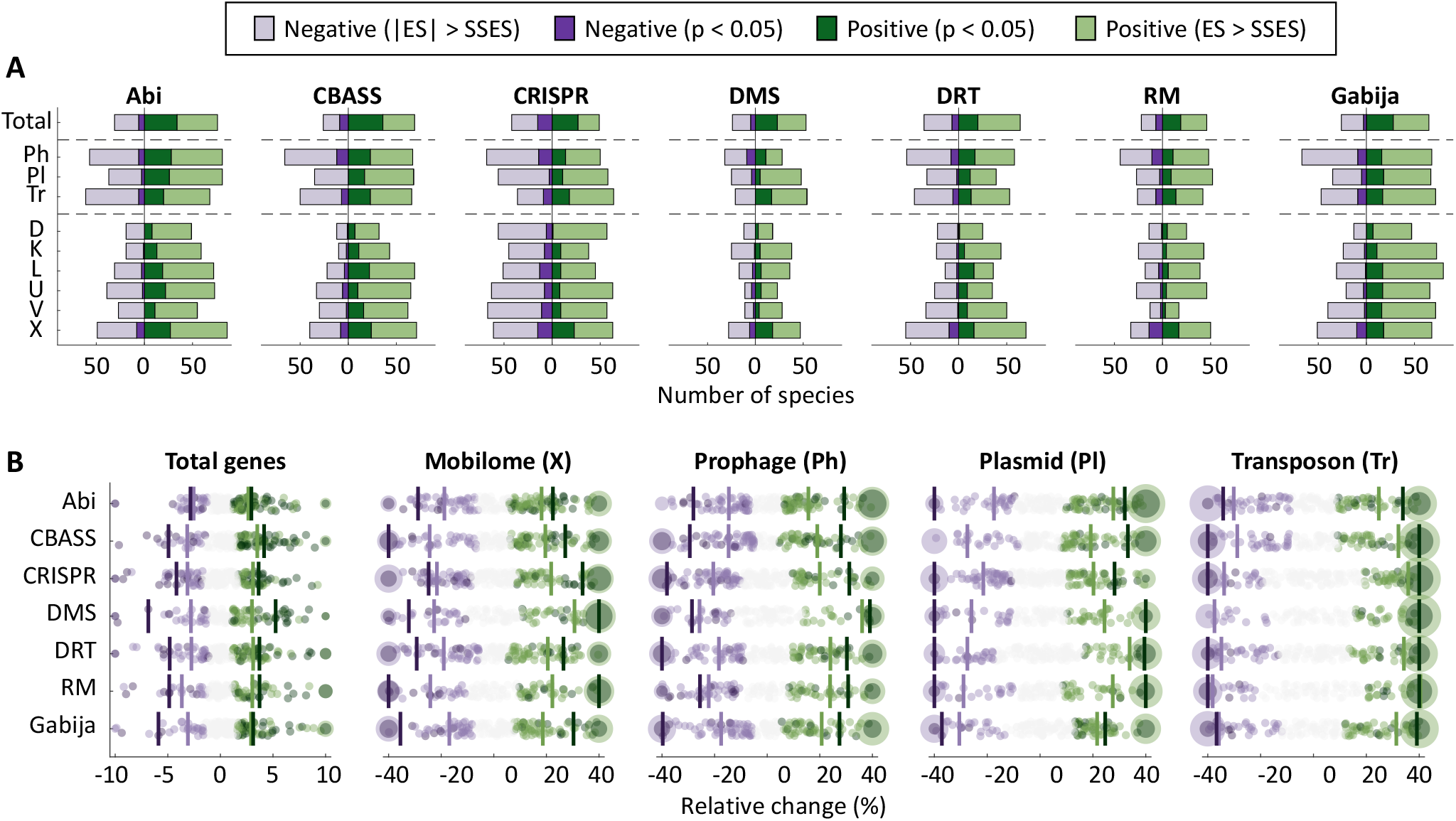
Association between 7 widespread defense systems, total number of genes, and MGE abundance. (a) Number of species displaying positive or negative associations in a phylogenetic generalized linear mixed effects model (see Methods) according to two different criteria: *p* < 0.05 and absolute effect size greater than the smallest significant effect size (|ES|>SSES), separately computed for each response variable and defense system. The top row (Total) indicates the association with the total number of genes. The association with MGE was calculated based on marker genes for prophages (Ph), plasmids (Pl), and transposons (Tr). Abbreviations of functional categories: X (mobilome), L (replication, recombination and repair), U (intracellular trafficking and secretion), K (transcription), D (cell cycle control, cell division and chromosome partitioning), and V (defense). (b) Effect sizes, measured as relative differences in gene and MGE abundances. Each point corresponds to one species. Species with values beyond the axis limits are collapsed in a single point with size proportional to the number of species. Vertical lines indicate the median over all the species that show positive or negative association according to the p-value and SSES criteria.

A more detailed analysis at the level of functional categories reveals that the presence of defense systems is most often associated with changes in the number of genes from COG categories X (mobilome), L (replication, recombination and repair), U (intracellular trafficking and secretion), K (transcription), D (cell cycle control, cell division and chromosome partitioning), and V (defense) (Fig. 1 and S1). Genes from these functional categories are typically present in MGE, suggesting that correlations (both positive and negative) between defense systems and genome size are primarily due to differences in the abundance of MGE. We confirmed that by masking genomic regions that correspond to MGE and rerunning the statistical analysis. As expected, 82% of the significant associations disappeared after masking MGE (Fig S2). Although a few significant associations persisted, a closer inspection revealed that those were often related to genes from prophages and other MGE (such as phage satellites) that had not been originally identified as such and remained unmasked.

The sign of the association between defense systems and MGE does not follow a clear taxonomic trend (Fig. 2, S3 and S4), with opposite signs sometimes found in closely related species (see, for example, the differences for CRISPR-Cas in *Phocaeicola vulgatus* and *P. dorei*). Moreover, the same species often display opposite trends for different defense systems. For example, in *Pseudomonas aeruginosa*, genomes with CRISPR-Cas contain fewer MGE, whereas genomes with CBASS, Gabija, and RM systems are significantly enriched in MGE. Negative associations between CRISPR-Cas and MGE are more abundant in the genus *Acinetobacter* (5 out of 8 species), the phylum *Bacteroidota* (negative association in 8 species, positive association in a single species) and the class *Clostridia*, the latter especially affecting prophages (negative association in 11 species, positive association in 2 species; Fig. S4). The genus *Acinetobacter* is also enriched in negative associations involving Abi (5 species) and CBASS (4 species). Furthermore, three almost non-overlapping groups of streptococci display negative associations for different defense systems. These encompass *S. pyogenes, S. gordonii, S. anginosus, S. mutans*, and *S. salivarius* in the case of CRISPR-Cas; *S. anginosus, S. oralis, S. intermedius*, and *S. suis* in the case of RM and DMS; and *S. pyogenes, S. dysgalactiae, S. equi*, and *S. uberis* in the case of Gabija.

**Figure 2:**
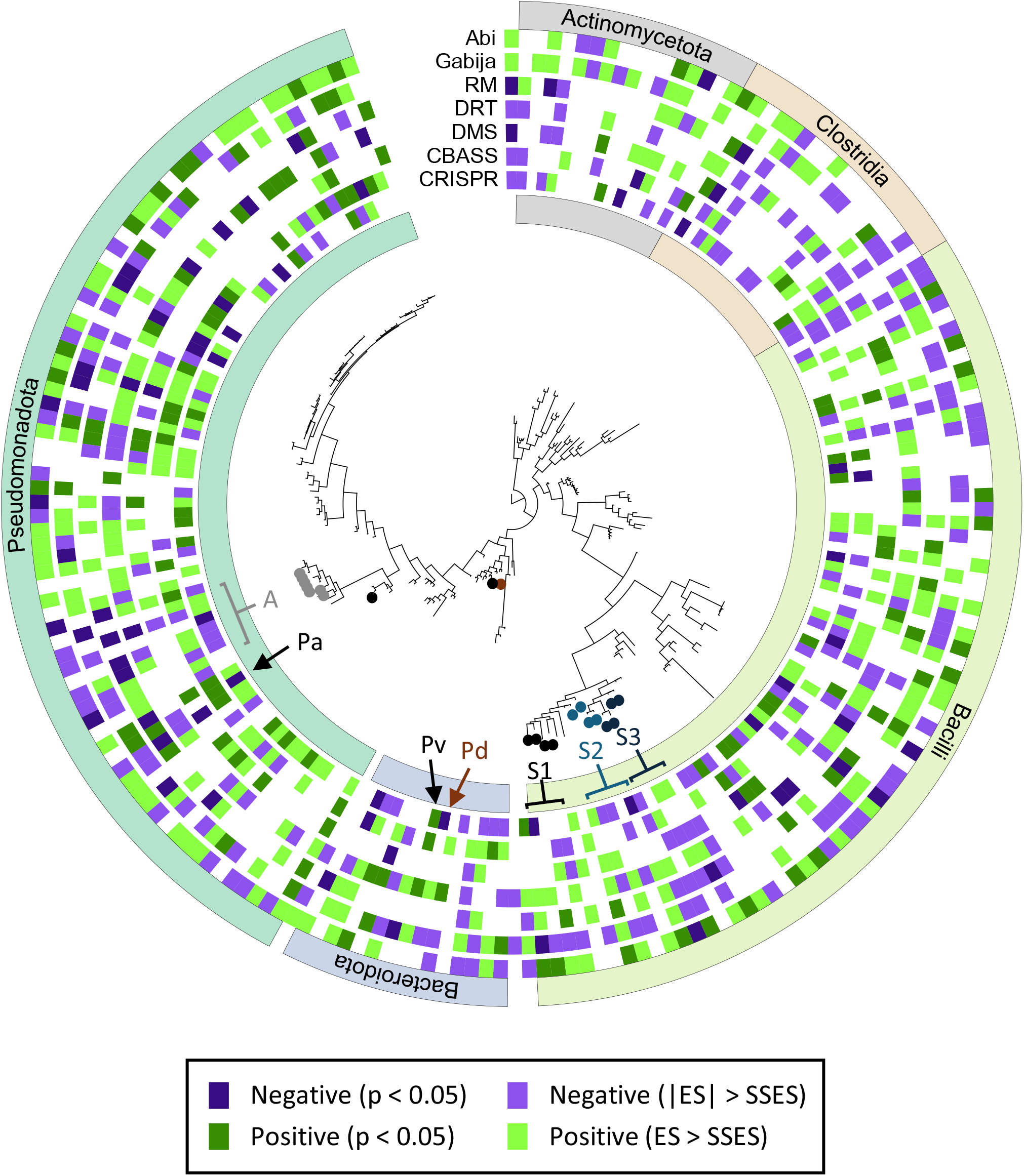
Taxonomic distribution of species displaying positive and negative associations between the presence of defense systems and the number of genes from the mobilome (based on COG annotations). Taxa discussed in the text are labeled: *A, Acinetobacter*; Pa, *Pseudomonas aeruginosa*; Pv, *Phocaeicola vulgatus*; Pd, *Phocaeicola dorei*; S1, *Streptococcus pyogenes, S. dysgalactiae, S. equi*, and *S. uberis; S2, S. gordonii, S. anginosus, S. mutans*, and *S. salivarius; S3, S. oralis, S. intermedius*, and *S. suis*. The classes Bacilli and Clostridia are the major components of the Genome Taxonomy Database phyla “Bacillota” and “Bacillota_A”. See figure S3 for a larger version including all species names.

The only archaeal species in the study (*Methanococcus maripaludis*) did not show any remarkable trend, other than a relatively strong (but not statistically significant) positive association between prophages and DMS/RM, and between transposons and Gabija, and a moderate negative association between prophages and Gabija.

### Associations between defense systems and MGE arise from differences in the rate of gene acquisition

To investigate if correlations between defense systems and MGE involve cross-strain differences in genome plasticity, we built high-resolution strain trees and identified clades that contain defense systems (DEF^+^). Then, we inferred the rates of gene gain and loss associated with different classes of MGE in DEF^+^ clades and in their respective sister groups lacking the defense system (DEF^-^) using the phylogenomic reconstruction tool GLOOME. We quantified the relative differences in gene gain and loss by dividing the rates observed in DEF^+^ clades by those from sister DEF^-^ clades. Finally, we compared the resulting DEF^+^/DEF^-^ gain and loss ratios between species in which MGE abundances and defense systems are positively and negatively correlated. We found that DEF^+^ and DEF^-^ clades often differ in their gain rates but not in their loss rates (Fig. 3). Specifically, in species in which defense systems are associated with increased MGE abundance, gene gain rates are typically higher in clades that contain the defense system. The opposite (that is, a reduction of gene gain rates in DEF^+^ clades) is observed in species in which defense systems are associated with lower MGE abundance. Among the latter, the biggest reductions in plasmid acquisition occur in association with RM, DRT, DMS and Abi, whereas significant drops in prophage gain are observed for DRT, CBASS and CRISPR-Cas. Taken together, these results confirm that correlations between defense systems and MGE arise from cross-strain differences in gene gain rather than loss, as expected given the role of MGE as facilitators of HGT.

**Figure 3:**
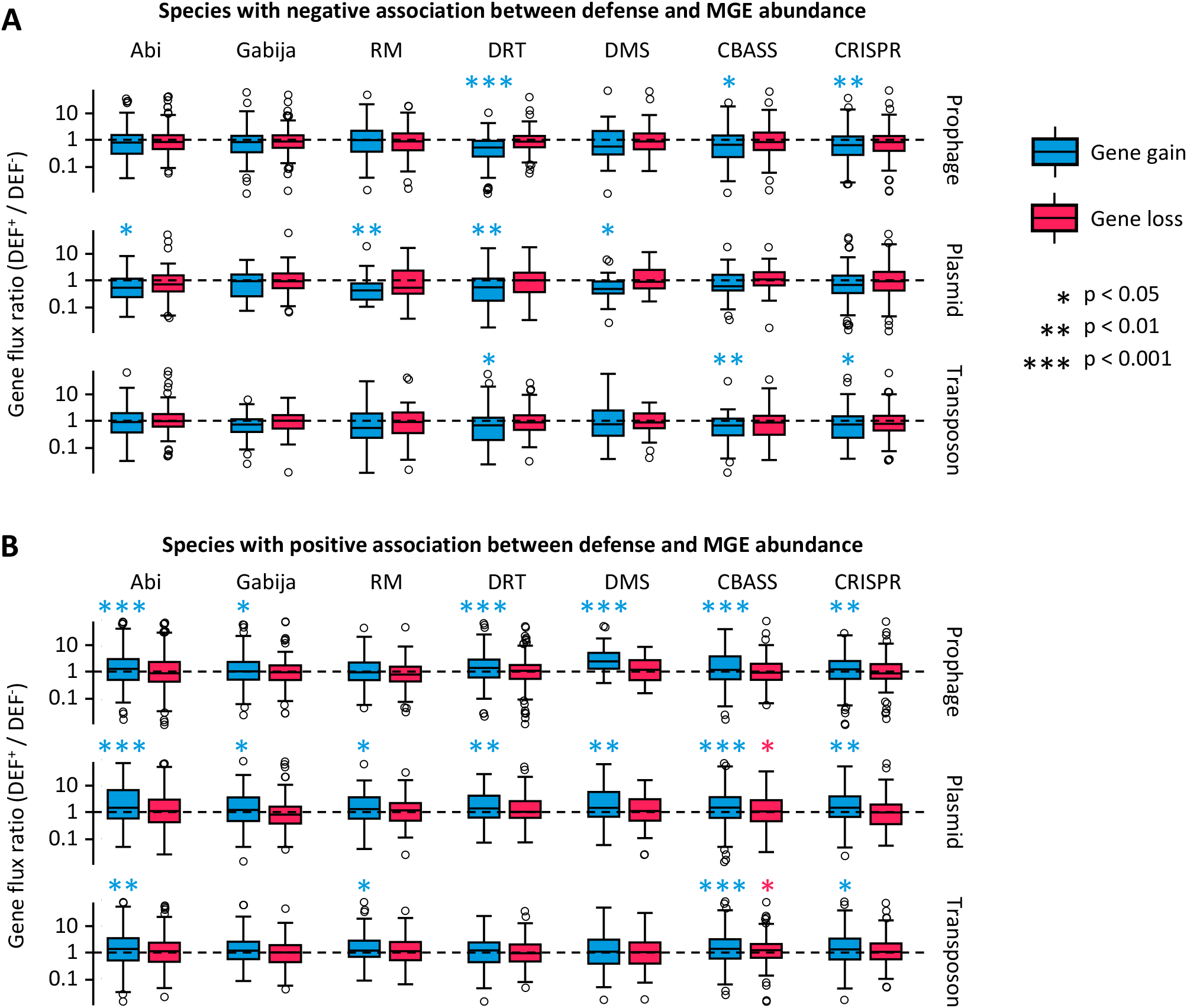
Relative differences in the rates of gene gain and loss between sister clades that do and do not harbor defense systems (DEF^+^ and DEF^-^, respectively). The boxplots represent the distribution of the DEF^+^/DEF^-^ ratio of gene gain (or loss) rates for each class of MGE, calculated for every pair of sister clades. Values greater (or smaller) than 1 indicate increased (or reduced) gene flux in lineages that contain the defense system. (a) Species that show a negative association between the defense system and the number of marker genes for each class of MGE (PGLMM with smallest significant effect size criterion). (b) Species that show a positive association between the defense system and the number of marker genes for each class of MGE. In the boxplots, the central line indicates the median, the box limits correspond to the 25 and 75 percentiles, and the whiskers extend to the largest and smallest values not classified as outliers. P-values are based on Wilcoxon test with log-ratio = 0 as null hypothesis.

### Timescales and linkage determine the sign of associations between defense systems and HGT

Because prokaryotic defense systems are often located within or adjacent to MGE, we hypothesized that positive associations between defense systems and MGE abundance could be explained, at least in part, by recent co-transfer events (henceforth, we term this the “linkage hypothesis”). Uder this hypothesis, positive associations would simply result from defense systems travelling together with MGE.

To test the linkage hypothesis, we first verified that defense systems tend to be co-transferred with MGE. We used ancestral reconstruction methods to identify the branches in which defense systems and MGE were gained and lost along strain trees (Table S5). We found that >95% of defense acquisition events occurred in branches in which MGE were also gained, and >90% of defense losses occurred in branches in which MGE were also lost (Fig. 4a). The random expectation given the rates of MGE gain and loss would be 71% and 63%, respectively (deviations with respect to these expectations are statistically significant with *p* < 10^−8^, binomial test). We also quantified the effect of MGE gain and loss on the per-branch probability of gaining or losing defense systems. The probability of acquiring a defense system is around 50-fold higher in branches in which MGE are gained than in other branches (Fig. 4b-c; Fisher exact test *p* < 10^−8^ for all defense systems). Similarly, the probability of losing a defense system is 10-to 50-fold higher if MGE are also lost in the same branch (Fig. 4b-c; Fisher exact test *p* < 10^−20^ for all defense systems). These trends are observed even in very short branches, spanning an evolutionary time of 10^−6^ substitutions per site in core genes (Fig. S5). To rule out the possibility that these associations were due to the presence of incomplete genomes, we separately considered terminal and internal tree branches. Because the dataset only includes complete and nearly complete genomes, the absence of a small number of missing genes, if relevant, would only affect the inference of gene gain and loss in terminal branches (those leading from the immediate ancestor to each incomplete genome). Despite some quantitative differences, strong co-acquisition (and co-loss) of MGE and defense occurs in both terminal and internal branches (Fig. S5), confirming that these associations are genuine.

**Figure 4:**
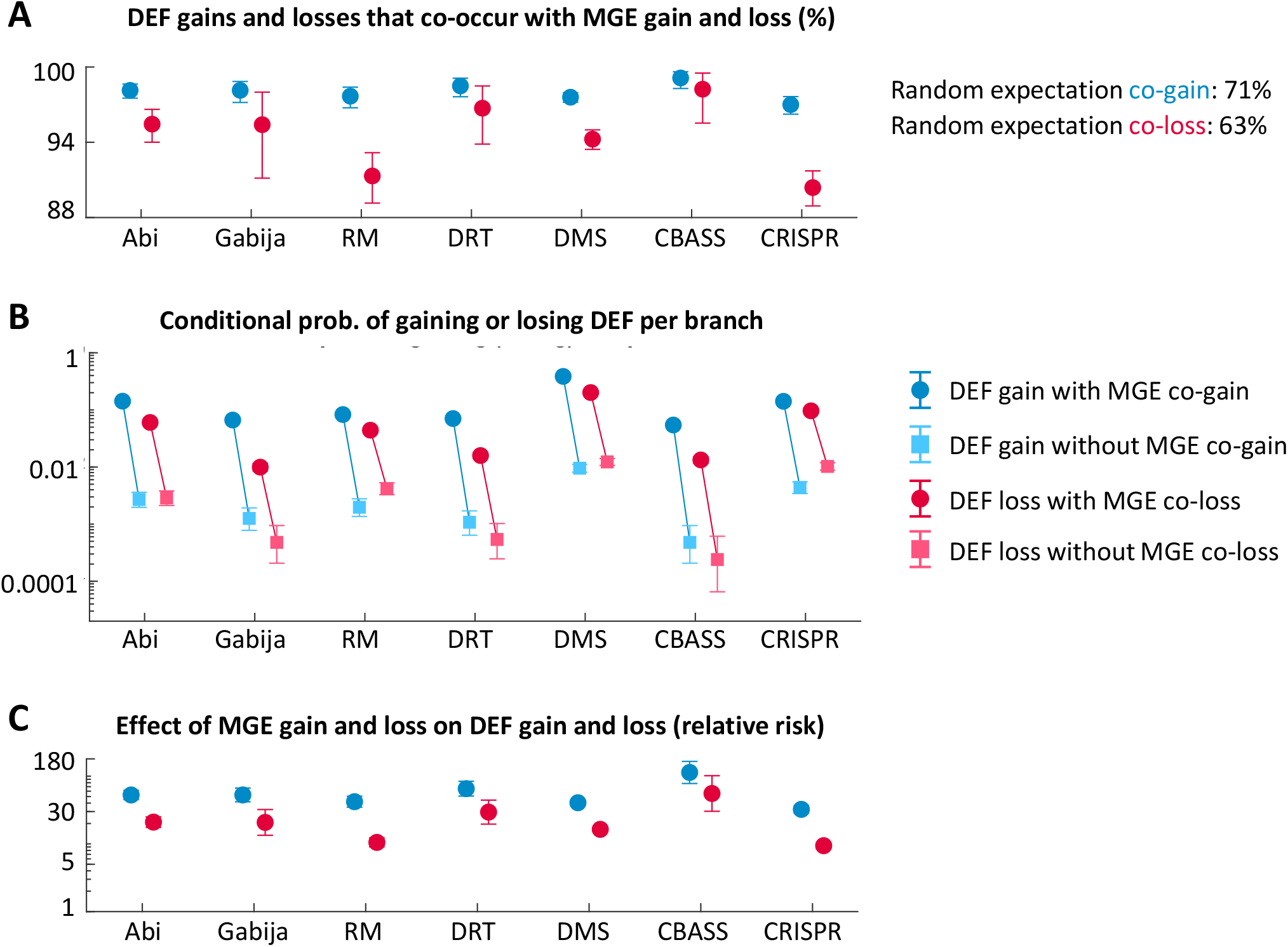
Co-occurrence of defense system gains and losses and MGE gains and losses along the phylogeny. (a) Percentage of DEF gains (and losses) that occur in the same branch as an MGE gain (or loss). (b) Conditional probability of gaining (or losing) a defense system provided that an MGE is also gained (or lost) in the same branch, compared to the conditional probabilities when an MGE is not acquired (or lost) in the same branch. (c) Effect of MGE gain (or loss) on the per branch probability to acquire (or loss) a defense system, measured as a risk ratio. Error bars in (a) and (b) correspond to 95% confidence intervals based on the binomial distribution. Error bars in (c) indicate the 95% confidence intervals for the relative risk (Morris and Gardner 1988).

Although co-gain of MGE and defense systems along the same tree branch does not necessarily imply a single event of joint gain, the strength of the association, the extremely short timespans, and the fact that similar trends are observed for gene losses strongly suggest that concurrent gain and loss of MGE and defense systems involve genetic linkage. (Alternative explanations in terms of strong selective pressure to quickly acquire defense mechanisms upon exposure to MGE and lose them once the MGE disappear might produce similar trends at the population level but not at the single-genome level, whereas episodic increases in the overall rate of HGT leading to separate but correlated acquisition of MGE and defense systems would not explain correlated losses.)

A major consequence of the co-transfer of MGE and defense systems is that the negative effects of defense systems on HGT should be easier to detect at longer timescales or under evolutionary conditions that weakened genetic linkage. To test that prediction, we studied how relative differences in the rates of gene gain between DEF^+^ and DEF^-^ clades depend on the depth of their last common ancestor in the strain tree (note that the depth of the last common ancestor serves as an upper limit for the time that the defense system has been retained in a lineage). As expected, positive outliers (with much higher gene gain rate in the DEF^+^ clade than in the DEF^-^ clade) almost invariably correspond to very recent lineages (less than 10^−5^ substitutions per bp in nearly universal core genes), indicating a very recent acquisition of the defense system (Fig. 5a and Table S6). In contrast, negative outliers (with much lower gene gain rate in the DEF^+^ clade than in the DEF^-^ clade) generally correspond to deeper lineages (at least 0.001 substitutions per bp). We quantified these trends by calculating the skewness of the distribution of DEF^+^/DEF^-^ log-transformed gain ratios in very recent and older lineages (Fig. 5b and Table S7). All the distributions show significant positive skewness in recent lineages and significant negative skewness in older lineages (all *p* < 0.001 except recent RM lineages, with *p* = 0.023; d’Agostino test for skewness). This quantitative analysis confirms that differences in timescale affect all defense systems, with positive associations between defense systems and gene gain being dominant in the short term and negative associations becoming more frequent in the long term.

**Figure 5:**
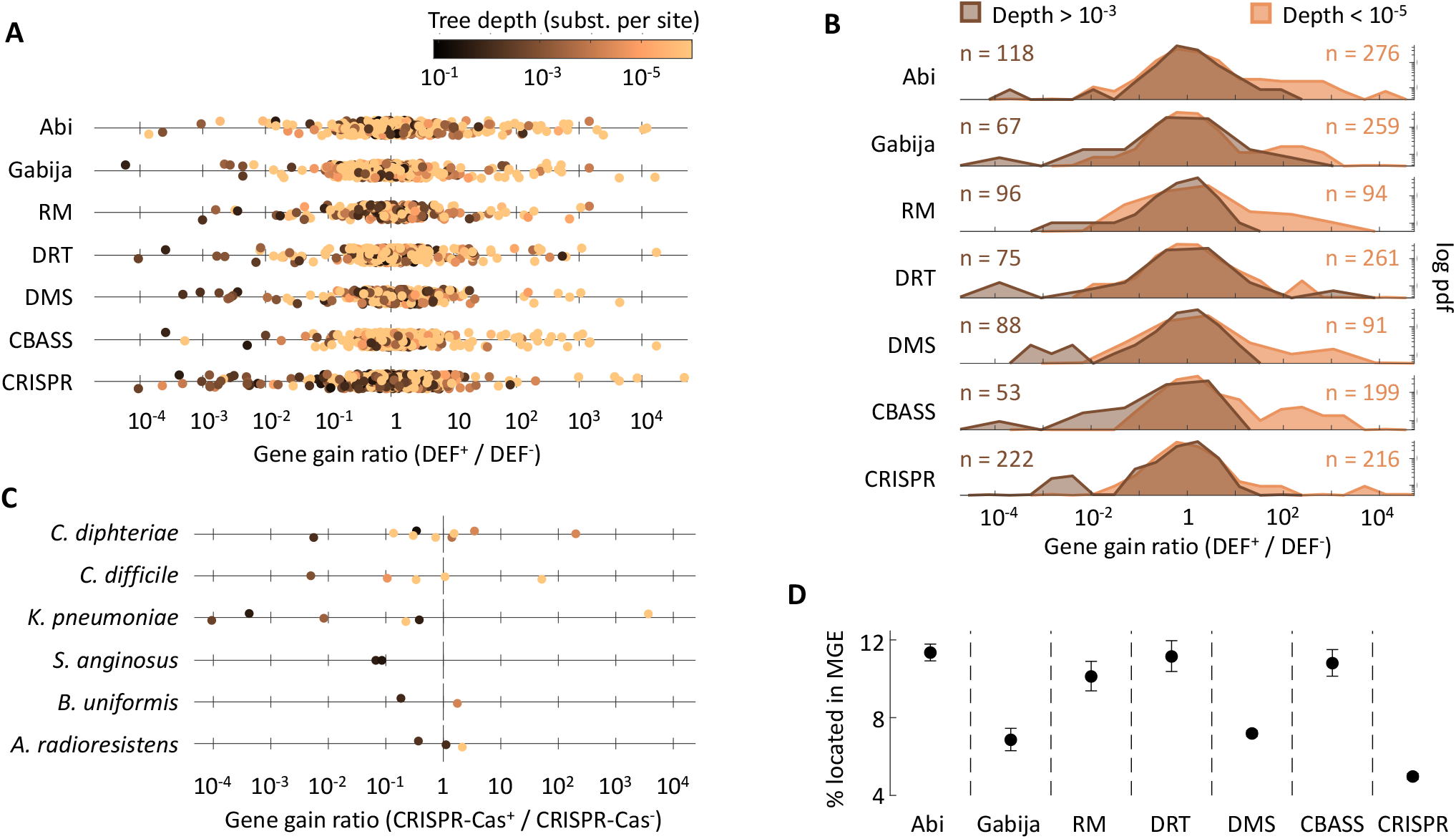
Association between defense systems and gene gain rates depends on the time scale. (a) Ratio of overall gene gain rates between sister clades that do and do not harbor defense systems (DEF^+^ and DEF^-^, respectively). Values greater (or smaller) than 1 indicate increased (or reduced) gene flux in lineages that contain the defense system. (b) Density distribution of the DEF^+^/DEF^-^ gain ratios, comparing clades that represent recent acquisitions of the defense system (depth < 10^−5^ substitutions per site in core genes) and clades that have retained the defense system for longer times (depth > 10^−3^ substitutions per site). The probability density function (y-axis) is represented in logarithmic scale to facilitate the visualization the tails. The skewness of all distributions is statistically significant (Table S7), with positive values for recent clades and negative values for deeper clades (c) Representative examples of gene gain ratios in shallow and deeper sister groups from the same species. (d) Percentage of genes from different defense systems located within known MGE. Whiskers represent 95% confidence intervals based on the binomial distribution.

A second prediction of the linkage hypothesis is that the net effect of defense systems on HGT is not only species specific, but also lineage specific. That is, defense systems may display positive, zero, or negative association with HGT in different lineages depending on when the defense system was acquired and how tight is the linkage to MGE. A more detailed study of gene gain rates in individual species confirms that the effect of CRISPR-Cas is, indeed, lineage-specific, with the same species encompassing recent CRISPR-Cas^+^ lineages that display increased gene gain rates and older CRISPR-Cas^+^ lineages with reduced gene gain rates (Fig. 5c). This observation, combined with the very recent acquisition of CRISPR-Cas in most lineages, could explain why many species show non-significant or positive associations between CRISPR-Cas and MGE abundances. Indeed, of the six representative species shown in Fig. 5c, only *Streptococcus anginosus* and *Acinetobacter radioresistens* show a net negative association between CRISPR-Cas and MGE abundance in the PGLMM analysis.

The findings described so far indicate that, of the seven defense systems considered in this study, CRISPR-Cas is the one that most often induces a net reduction in HGT rates. According to the linkage hypothesis, this could be due to a comparatively weaker physical association between CRISPR-Cas and MGE. To evaluate that possibility, we calculated the percentage of genes from CRISPR-Cas located inside MGE and compared that to other defense systems (Fig. 5d). Consistent with the linkage hypothesis, CRISPR-Cas is the defense system that is least often encoded by MGE (4.97% vs 8.12%, *p* < 10^−8^, chi-squared test), followed by Gabija and DMS.

### Anti-CRISPR proteins modulate associations between CRISPR-Cas and HGT

Anti-CRISPR proteins (Acr) have the potential to suppress the possible negative effect of CRISPR-Cas on MGE-driven gene transfer (Mahendra et al. 2020). This, in turn, could contribute to explaining the sign of the association between CRISPR-Cas and MGE in different lineages. To assess that possibility, we searched all the genomes in the dataset for known Acr, finding 16,058 proteins. Then, we compared the prevalence of Acr in species in which the presence of CRISPR-Cas is positively or negatively correlated with the MGE content, separately considering genomes that do and do not harbor CRISPR-Cas. Because Acr are generally encoded by MGE (Pinilla-Redondo et al. 2020) and the prevalence of MGE systematically varies across groups acting as a confounding factor, we restricted our comparisons to genomes that contain at least one prophage. Our results indicate that genomes with and without CRISPR-Cas differ in the prevalence of Acr (Fig. S6). More importantly, these differences have opposite directions in species in which CRISPR-Cas is positively and negatively associated with MGE abundance. In the former, Acr are more prevalent in genomes that contain CRISPR-Cas (p = 0.0012, chi-squared test). In the latter, Acr are less prevalent in genomes with CRISPR-Cas (*p* < 10^−6^, chi-squared test). In both groups, the prevalence of Acr in genomes without CRISPR-Cas is similar. These results suggest that negative associations between CRISPR-Cas and HGT could be dependent on (or at least facilitated by) low prevalence of Acr in the genome.

## Discussion

Defense systems could have a significant impact on microbial evolution by effectively blocking the transfer of MGE, reducing gene flow and limiting the spread of accessory genes. However, because defense systems are often carried by MGE and these are the main vehicles of HGT, a net positive association between defense systems and gene exchange cannot be ruled out a priori. We assessed the relative weight of these two opposite scenarios by quantifying the association between defense systems, MGE abundance, and gene acquisition rates in a phylogeny-aware comparative study of 197 prokaryotic species.

Our results shed light on previous, apparently contradictory findings concerning the effect of CRISPR-Cas on genome evolution and diversification (Gophna et al. 2015; O’Meara and Nunney 2019; Shehreen et al. 2019; Wheatley and MacLean 2021; Pursey et al. 2022). A pioneering study conducted in 2015 found no evidence to support an overall association between CRISPR-Cas activity and reduced gene acquisition via HGT at evolutionary time scales (Gophna et al. 2015). Such lack of association was explained by several factors, including the high mobility of CRISPR-Cas systems, that limits their long-term impact on host genomes, and the possibility that HGT is mediated by MGE that escape (or are not targeted by) CRISPR-Cas immunity.

Our analyses support the general conclusion that CRISPR-Cas and other defense systems have little overall impact on HGT in most bacterial species. Specifically, differences in the rates of gene acquisition in lineages that do and do not harbor defense systems are centered around zero. That said, we identified significant opposite trends at very short and intermediate evolutionary time scales: positive associations between defense systems and HGT are more frequent at very short time scales, whereas negative associations become dominant at longer time scales. These opposite trends suggest that the actual effects of defense systems on gene exchange may be obscured by recent co-transfer events involving MGE. As a result, the possible negative effects of defense systems on HGT only become detectable if the defense system is maintained for long enough periods of time.

Besides this general picture, we identified some species in which the presence of defense systems (especially CRISPR-Cas) significantly correlates with smaller genome sizes and MGE abundances. Some of those associations had been previously described in *P. aeruginosa* and *Klebsiella pneumoniae* (Wheatley and MacLean 2021; Botelho et al. 2023). And yet, these species represent special cases rather than the rule, even in the context of host-associated bacteria. In fact, our results underline that the association between defense systems and HGT is strongly system- and lineage-dependent. This conclusion confirms and extends previous findings that showed that the impact of CRISPR-Cas on the spread of antibiotic resistance is highly variable across species and its sign cannot be easily explained by simple ecological, environmental, or genomic variables (Shehreen et al. 2019).

Among the 7 defense systems included in this study, CRISPR-Cas stands out for being the most recurrently associated with reduced genome sizes and lower MGE (especially prophage) abundances. In contrast, DMS, Abi, DRT, and CBASS are more often associated with higher numbers of MGE and accessory genes. This finding is fully consistent with a recent study that compared 73 defense systems in 12 bacterial species (Kogay et al. 2024). We propose that what makes CRISPR-Cas systems different is their weaker (though still substantial) linkage with MGE. Compared to fully functional CRISPR-Cas systems, other defense systems like Abi, Gabija, CBASS, DMS, and RM are more frequently located within or next to MGE (Makarova et al. 2011; Benler et al. 2021; Rousset et al. 2022; Botelho 2023) and may have alternative functions related to MGE propagation. For example, Abi systems have been identified in PICIs as accessory genes that facilitate their parasitic lifecycle (Ibarra-Chavez et al. 2021). The narrow specificity of some defense systems may be another reason why those systems do not significantly interfere with HGT. For instance, the GmrSD type IV RM system selectively targets phages with glucosylated hydroxymethylcytosine (Bair and Black 2007) and the Thoeris defense system only appears to be effective against myoviruses (Doron et al. 2018). This caveat extends to any defense system based on epigenetic modifications, such as RM and DMS, whose overall effect on HGT depend on the repertoire of epigenetic markers in the host and MGE populations (Oliveira et al. 2016). Although selective targeting is often viewed as an outcome of phage-host coevolution, it is tempting to speculate that it could have been evolutionarily favored by the need to fight harmful genetic parasites while maintaining sufficiently high rates of HGT to prevent population-level gene loss (Iranzo et al. 2016).

Defense systems sometimes exhibit synergistic interactions (Dupuis et al. 2013; Wu et al. 2024), which could contribute to the heterogeneity of effects reported in this study. The vast number of potential interactions and the limited availability of high-quality genomes made it unfeasible to systematically account for the effect of interactions with the methods developed in this work. Determining if and to what extent synergy among defense systems affects HGT remains a subject for future investigation, possibly focused on a small set of experimentally validated interactions in highly sequenced species. Another open question concerns which levels of detail, both taxonomic and functional, best capture the effect of defense systems on HGT. From a taxonomic perspective, working at or below the species level is a natural choice because species represent genetically cohesive units (Bobay and Ochman 2017; Konstantinidis 2023; Conrad et al. 2024) and, as a result, uncontrolled confounding factors are less likely to affect within-species than cross-species comparisons. In contrast, more complex multi-level approaches would be required to detect trends at higher taxonomic ranks. From a functional perspective, we grouped defense systems based on their mechanism of action, under the assumption that functionally similar systems produce similar effects on HGT. Though reasonable, this grouping criterion may not be optimal in systems in which subtypes markedly differ in their eco-evolutionary dynamics and linkage with MGE. Moreover, fine-grain dissection of highly abundant systems, such as RM and DMS, could help improve the sensitivity of statistical tests by producing more balanced sets of strains with and without the subtypes of interest.

All in all, we showed that some defense systems, especially CRISPR-Cas, can significantly reduce HGT, although the effect is often masked by the fact that these systems travel together with MGE. Beyond possible functional connections, the linkage between defense systems and MGE is an inevitable consequence of the arms race between parasites and hosts. Because defense systems are costly and their efficacy drops as parasites evolve, they are subject to rapid turnover and depend on HGT for long-term persistence in microbial populations (van Houte et al. 2016; Iranzo et al. 2017; Koonin et al. 2017; Puigbo et al. 2017). As a result, it is extremely challenging to disentangle the impact of defense systems on gene flow from the causes that lead to their presence or absence, especially at short evolutionary time scales. As more and more genomic data become available, we expect that future research will overcome this challenge by quantifying the linkage between defense systems and MGE, developing more realistic null models, and testing the role of defense systems on microbial adaptation at different time scales.

## Methods

### Genome collection and identification of defense systems

We parsed the Genome Taxonomy Database (GTDB, https://gtdb.ecogenomic.org) release 202 (Parks et al. 2020) to identify all high-quality genomes (according to the MIMAG criteria (Bowers et al. 2017)) with completeness >99%, contamination <1%, and contig count <500. The 82,595 genomes that passed these filters were downloaded from the NCBI FTP site (https://ftp.ncbi.nlm.nih.gov). CRISPR-Cas systems were identified with CRISPRCasTyper v1.2.4 (Russel et al. 2020) using default parameters. We classified a genome as CRISPR-Cas^+^ if it contains at least one high-confidence Cas operon and CRISPR-Cas^-^ otherwise. Only the species with >10 genomes and at least 5 CRISPR-Cas^+^ genomes were further considered. To reduce the computational cost, we only considered a maximum of 500 genomes per species. Species with >500 genomes were randomly subsampled to keep at most 350 CRISPR-Cas^+^ and 150 CRISPR-Cas^-^ genomes. Other defense systems were identified with Padloc v1.1.0 (db v1.4.0) (Payne et al. 2022). Our analysis focused on the most prevalent defense systems: restriction-modification (RM), DMS, Abi, CRISPR-Cas, Gabija, DRT, and CBASS.

After applying these criteria, 19,323 genomes belonging to 196 bacterial and 1 archaeal species (sensu GTDB) were included in the analysis (Tables S1 and S2). Of those, 2,964 correspond to complete genomes and the rest to high-quality, nearly complete ones.

### Gene prediction and annotation

Open reading frames (ORF) were predicted with Prodigal v2.6.3, using codon table 11 (prokaryotic genetic code) and “single” mode, as recommended for finished and draft quality genomes (Hyatt et al. 2010). Orthologous ORF were then separately clustered for each species with Roary v3.13.0 (Page et al. 2015) setting an 80% identity threshold for initial clustering followed by synteny-based refinement (options ‘-t 11 -i 80’). The resulting gene clusters were functionally annotated by selecting a representative sequence, arbitrarily chosen among those with length between 0.95 and 1.05 times the average length of all sequences in the cluster. Representative sequences were functionally annotated by mapping them to in-house profiles of the Clusters of Orthologous Genes (COG) database (2020 release) (Galperin et al. 2021) with HMMER v3.1b2 (e-value < 0.001) (http://hmmer.org). The 26 major prokaryotic functional categories defined in the COG database were assigned to the annotated genes. Some functional categories (A, RNA processing and modification; B, chromatin structure and dynamics; W, extracellular structures; T, signal transduction; and Z, cytoskeleton) were excluded since they rarely or never occur in prokaryotic genomes. The case-insensitive keywords “phage”, “plasmid” and “transpos^*^” in the COG gene annotations were used to identify genes associated with prophages, plasmids, and transposons, respectively, and the resulting gene lists were manually curated to minimize false assignments. We based our statistical analyses on marker gene counts rather than full MGE counts because the latter are more susceptible to technical artifacts (e.g., different heuristics to deal with nested MGE will affect the number of MGE, but not the number of MGE marker genes). Gene counts per genome and functional category are listed in Table S8. Anti-defense proteins, including anti-CRISPR, were identified by running HMMER v3.1b2 (e-value < 10^−10^) against the dbAPIS database (Yan et al. 2024).

### Identification of genomic regions containing mobile genetic elements

Prophages were detected with Phispy v4.2.21 using default options (Akhter et al. 2012). Short transposons were identified based on the presence of isolated or paired genes annotated as transposases. In the latter case, we allowed for up to one additional gene between two transposon-related genes to account for the genetic architecture of some insertion sequences (Gomez et al. 2014). Due to the inherent difficulty to discriminate among plasmids, ICE, and IME (Botelho et al. 2023), we restricted the search for these elements to complete genomes. Plasmids were identified as extra-chromosomal replicons that contain the “plasmid” label in their NCBI description lines. After removing those replicons, ICEfinder (Liu et al. 2019) was used to identify ICE and IME. Finally, we ran Phispy v4.2.21 with option ‘-phage_genes 0’ to identify any other MGE, integrons, pathogenicity islands, and fragments of MGE that could have been missed by the previous approaches. The MGE identified through these approaches were masked from complete genomes to produce the gene counts in Table S9 and the results shown in Fig S2.

### Species trees

Phylogenetic trees were separately built for each species based on the set of 120 prokaryotic marker genes (122 in the case of Archaea) proposed by the GTDB r202. For each species, only those marker genes with prevalence >80% were used for phylogenetic reconstruction. We aligned the amino acid sequences of each marker gene with mafft-linsi (L-INS-I algorithm, default options, MAFFT v7.475) (Katoh and Standley 2013) and back-translated the amino acid alignments to nucleotide alignments with pal2nal.pl v14 (Suyama et al. 2006) using codon table 11. After concatenating all nucleotide alignments, we built preliminary trees with FastTree v2.1.10 (options ‘-gtr -nt -gamma -nosupport - mlacc 2 -slownni’) (Price et al. 2010). The tree topologies produced by FastTree were subsequently provided to RaxML v8.2.12 (Stamatakis 2014) for branch length optimization (raxmlHPC with options ‘-f e -m GTRGAMMA’). The final trees are included in Supplementary File S1.

To visualize trends across species (Fig. 2, S3, and S4), we used the online tool iTOL (Letunic and Bork 2021) and the multispecies tree from GTDB r202.

### Phylogenetic generalized linear mixed models (PGLMM)

For each genome in the dataset, we collected the following response variables: the total number of genes, the number of genes belonging to each functional category, and the number of genes associated with prophages, plasmids, and transposons (Table S8). Genes that belong to the 7 defense systems of interest were excluded when computing these values. Then, for each response variable, we fitted a PGLMM with Poisson distribution and canonical link function, using the presence or absence of each defense system as predictors and the species trees as guides to generate the covariance matrix.

For each species, the PGLMM assumes that the response variable, *Y*_*i*_, follows a Poisson distribution with mean *μ*_*i*_, that is, *Y*_*i*_∼ Poisson (*μ*_*i*_). The expected gene abundance *μ*_*i*_, is modeled as log *μ*_*i*_= *β*_0_ +∑ *β*_*j*_*X*_*ij*_+ *ϵ*_*i*_, where is *β*_0_ the intercept, *β*_*j*_ is the coefficient associated with the defense system *j*, and *X*_*ij*_ ∈ {0,1} denotes the absence or presence of defense system *j* in genome *i*. The random effects *ϵ*_*i*_ follow a multivariate normal distribution, 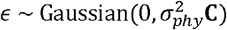 where 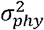 is the strength of the phylogenetic signal and **C** is a covariance matrix derived from the phylogenetic tree under the assumption of Brownian motion evolution.

To extend the PGLMM to multiple species, we considered that the effect of defense systems on gene content can be species-dependent (this was a reasonable assumption *a priori* and later confirmed by the analysis). To account for that, the multi-species model must include an interaction term “defense × species”, whose coefficients are relevant per se, and, accordingly, modeled as fixed effects. Moreover, because HGT rates are highly variable among species (Puigbò et al. 2014; Iranzo et al. 2019), it makes little sense to extend the phylogenetic correction beyond single species or assume that the strength of the phylogenetic signal is the same for all species. These considerations are captured by a phylogenetic covariance matrix with block-diagonal structure (one block per species), and species-wise values of the phylogenetic coefficient (one for each block of the phylogenetic covariance matrix). In practice, fitting a multi-species model with these specifications is formally equivalent to fitting independent models for each species. The latter approach, that we adopted, has the advantage of being more suitable for parallelization and requiring fewer computational resources. Thus, for each species and response variable, we fitted a PGLMM with the function pglmm_compare(response_variable ∼ Abi + CBASS + CRISPR + DMS + DRT + Gabija + RM, family = “poisson”, data = SpData, phy = SpTree) from the R package phyr v1.1.0 (Li et al. 2020). In the formula, Abi, CBASS, CRISPR, and so on, are binary variables representing the presence (1) or absence (0) of each defense system. Table S4 presents the coefficients, p-values, and goodness of fit of the model.

The model described above includes nine coefficients (intercept, seven defense systems, and the phylogenetic signal), which could lead to overfitting in species in which the number of available genomes is limited. As an alternative, we also fitted seven separate PGLMM, one for each defense system, involving a single predictor and the phylogenetic random effect (Table S3). This approach does not account for correlations among defense systems, but it has the advantage of not being affected by overfitting. The figures in the manuscript are based on this second set of models, although, in practice, both approaches produce very similar quantitative results.

For each class of PGLMM, we also fitted non-phylogenetic models in which the covariance matrix **C** of the random effect was replaced by the identity matrix. The PGLMM were compared to their non-phylogenetic counterparts using the conditional Akaike Information Criterion (cAIC) as previously described (Greven and Kneib 2010; Säfken et al. 2021). Based on the cAIC, PGLMM performed better than non-phylogenetic models in 65% of the species-response-defense triplets and were close to non-phylogenetic models (ΔcAIC < 2) in another 25% of the triplets (in most of those cases, the strength of the phylogenetic signal was close to zero, which made both phylogenetic and non-phylogenetic models equivalent).

To account for the possibility that other (less abundant) defense systems could explain part of the variability in the results, we explored a more complex set of models that included the total number of other defense systems as an additional predictor. These models generally performed worse than their simpler variants (ΔcAIC > 0 in 83% of the species) and were not considered for further analysis.

Large differences in sample size among species and defense systems translate into unequal precision in the estimation of the model coefficients. To deal with that limitation, we used two different criteria to identify species in which the presence of a defense system is positively or negatively associated with gene numbers. For one option, we adopted a classical criterion of statistical significance (*p* < 0.05) for the predictor variable in the PGLMM. For the other option, we applied an alternative criterion based on effect size, aimed at comparing species with different sample sizes in which p-values are not commensurable. Specifically, for each variable and defense system, we jointly considered the PGLMM of all the species and determined the smallest effect size (in absolute value) that reached statistical significance (*p* < 0.05) in any species. Then we used the smallest significant effect size (SSES) as a threshold to classify associations as positive, negative, or null.

### Inference of gene gain and loss

Gene gains and losses at each branch of each species tree were estimated with Gloome (Cohen and Pupko 2010), using as inputs the gene presence/absence matrices previously generated by Roary and the species trees. The parameter configuration file was set to optimize the likelihood of the observed phyletic profiles under a genome evolution model with 4 categories of gamma-distributed gain and loss rates and stationary frequencies at the root.

### Comparison of gene gain and loss rates between DEF^+^ and DEF^-^ clades

For each species tree, we defined DEF^+^ clades as the narrowest possible clades such that at least 80% of the leaves contain the defense system of interest. Candidate DEF^-^ clades were defined in an analogous way, referring to leaves without the defense system. Next, we identified pairs of DEF^+^ and DEF^-^ clades that constitute sister groups. Sister DEF^+/-^ pairs were excluded if both clades contained a single genome. For each clade in a valid pair, we computed the overall gene gain and loss rates as the expected number of gene gains (or losses) in that clade divided by the total branch length. Gain and loss rates for different functional categories and MGE we calculated in an analogous way but restricting the sum of gene gains and losses to the genes of interest. When calculating overall and category-wise gain and loss rates, we did not take into account the contribution of species-wise singletons (genes without homologs in other genomes of the same species), as they may represent false gene predictions or genes that are replaced at unusually high rates (Wolf et al. 2016). To account for the non-negative nature of gain and loss rates and their heavy-tailed distributions, comparisons between DEF^+^ and DEF^-^ sister branches were done based on log-transformed rate estimates. To calculate the skewness of the distributions and their statistical significance, we use the method proposed by D’Agostino (D’Agostino et al. 1990) as implemented by the scipy.stats.skewtest() function in Python (Virtanen et al. 2020).

## Data access

All the data generated or analyzed during this study are included in this published article and its supplementary information files.

## Competing interests

The authors declare no competing financial interests. Acknowledgements Y.L. is supported by China Scholarship Council (No.202008440425). J.B. is supported by the Maria Zambrano grant of the Spanish Ministry of Universities (Grant No. UP2021-035), and the Severo Ochoa Program for Centres of Excellence in R&D of the Agencia Estatal de Investigación of Spain (Grant No. CEX2020-000999-S (2022–2025) to the CBGP). J.I is supported by the Ramón y Cajal Programme of the Spanish Ministry of Science (Grant No. RYC-2017–22524); the Agencia Estatal de Investigación of Spain (Grant Nos. PID2019-106618GA-I00 and CNS2023-145430), the Severo Ochoa Programme for Centres of Excellence in R&D of the Agencia Estatal de Investigación of Spain (Grant No. SEV-2016–0672 (2017–2021) to the CBGP); and the Comunidad de Madrid (through the call Research Grants for Young Investigators from Universidad Politécnica de Madrid, Grant No. M190020074JIIS).

We thank Jorge Calle-Espinosa for helpful discussions.

