## Supplemental Figure S1 for "Timescale and genetic linkage explain the variable impact of defense systems on horizontal gene transfer"

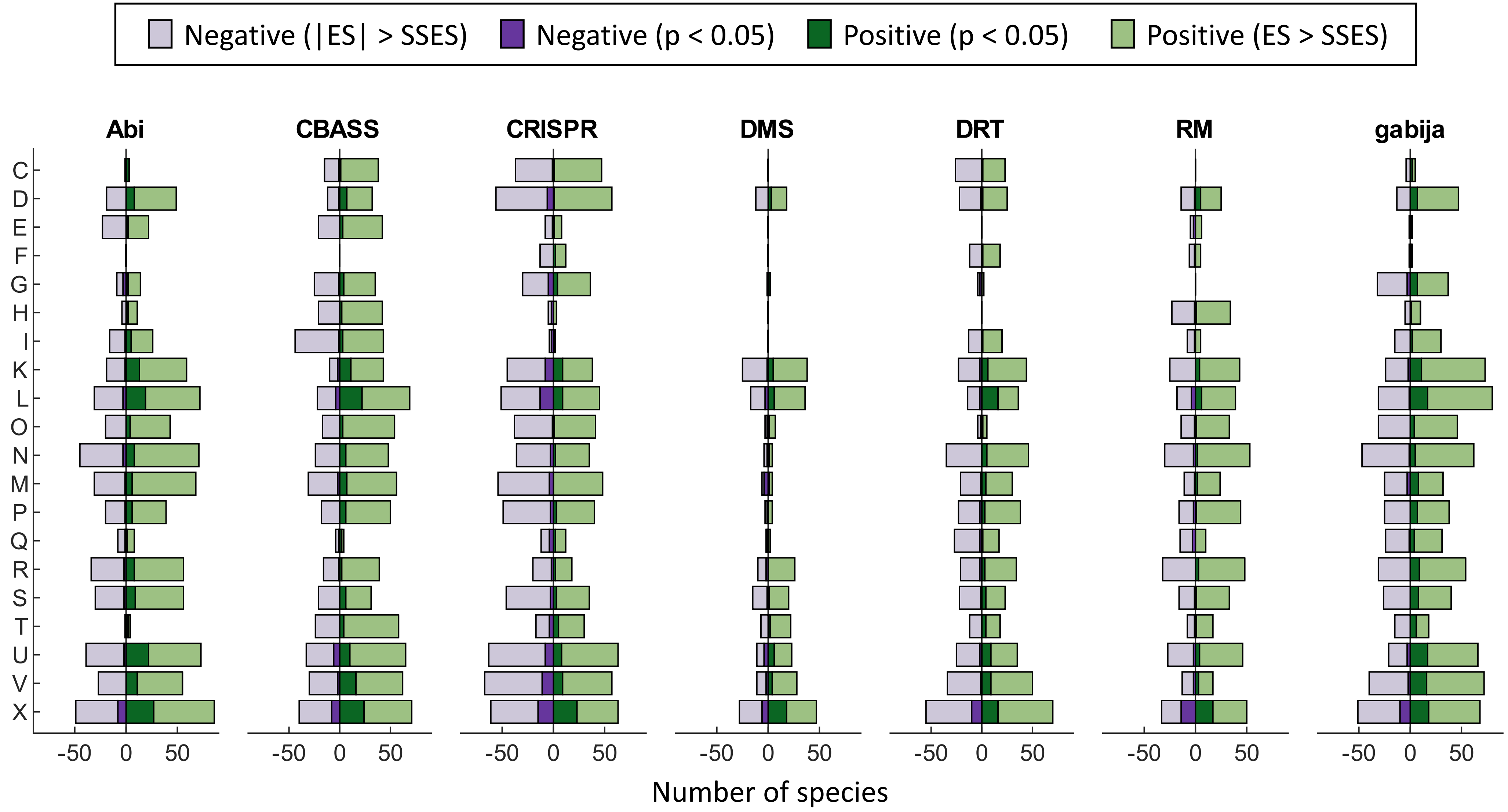


**Figure S1:** Association between 7 widespread defense systems and the number of genes from different functional categories (based on COG annotations). The length of the bars indicates the number of species displaying positive or negative associations in a phylogenetic generalized linear mixed effects model (see Methods) according to two alternative criteria: $p < 0.05$ and absolute effect size greater than the smallest significant effect size (|ES|>SSES), separately computed for each response variable and defense system. Abbreviations of functional categories, C: energy production and conversion; D: cell cycle control, cell division, chromosome partitioning; E: amino acid transport and metabolism; F: nucleotide transport and metabolism; G: carbohydrate transport and metabolism; H: coenzyme transport and metabolism; I: lipid transport and metabolism; J: translation, ribosomal structure and biogenesis; K: transcription; L: replication, recombination and repair; M: cell wall/membrane/envelope biogenesis; N: cell motility; O: posttranslational modification, protein turnover, chaperones; P: inorganic ion transport and metabolism; Q: secondary metabolites biosynthesis, transport and catabolism; R: general function prediction only; S: function unknown; T: signal transduction mechanisms; U: intracellular trafficking, secretion, and vesicular transport; V: defense mechanisms; X: mobilome: prophages, transposons.
