## Supplemental Figure S2 for "Timescale and genetic linkage explain the variable impact of defense systems on horizontal gene transfer"

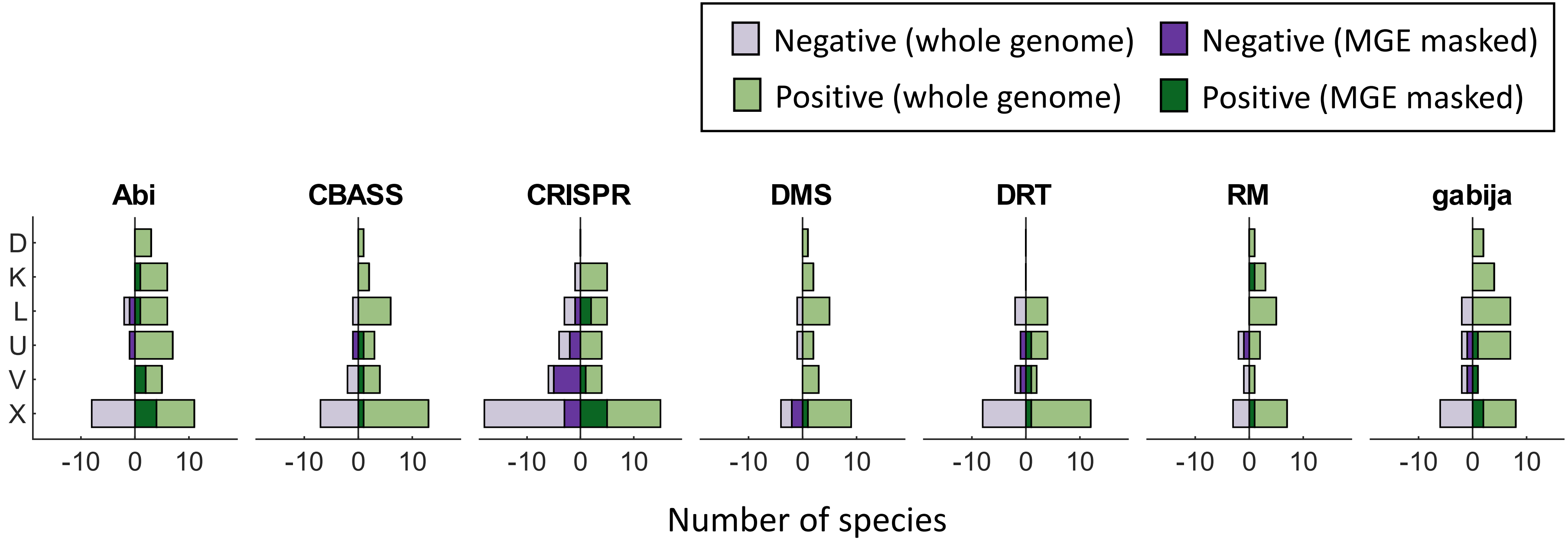


**Figure S2:** Association between 7 widespread defense systems and the number of genes from different functional categories before and after masking MGE from complete genomes. The length of the bars indicates the number of species displaying significant ($p < 0.05$) positive or negative associations in a phylogenetic generalized linear mixed effects model. Abbreviations of functional categories: X (mobilome), L (replication, recombination and repair), U (intracellular trafficking and secretion), K (transcription), D (cell cycle control, cell division and chromosome partitioning), and V (defense).
