## Supplemental Figure S3 for "Timescale and genetic linkage explain the variable impact of defense systems on horizontal gene transfer"

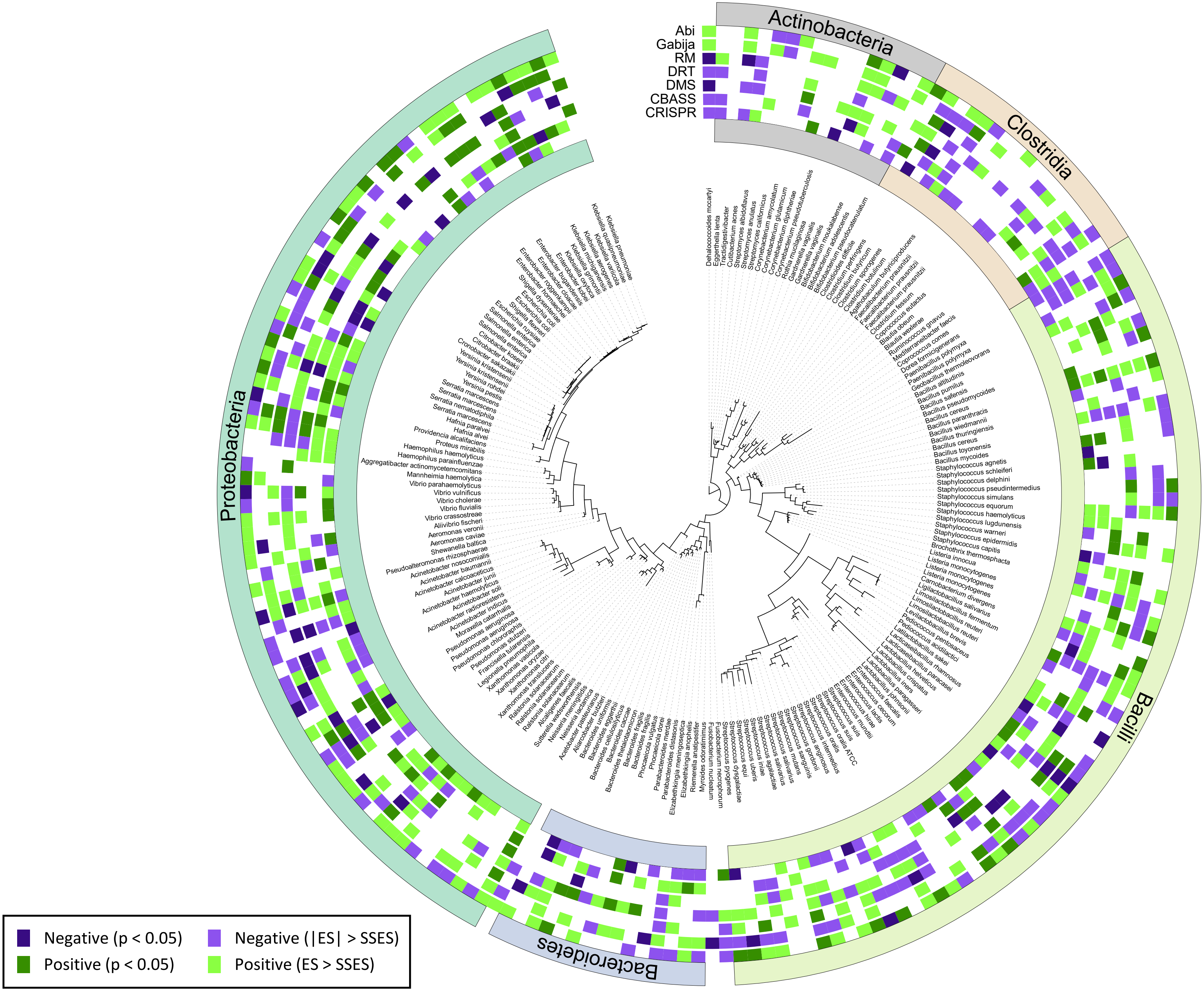


**Figure S3:** Taxonomic distribution of species displaying positive or negative associations between the presence of defense systems and the number of genes from the mobilome (based on COG annotations).
