## Supplemental Figure S4 for "Timescale and genetic linkage explain the variable impact of defense systems on horizontal gene transfer"

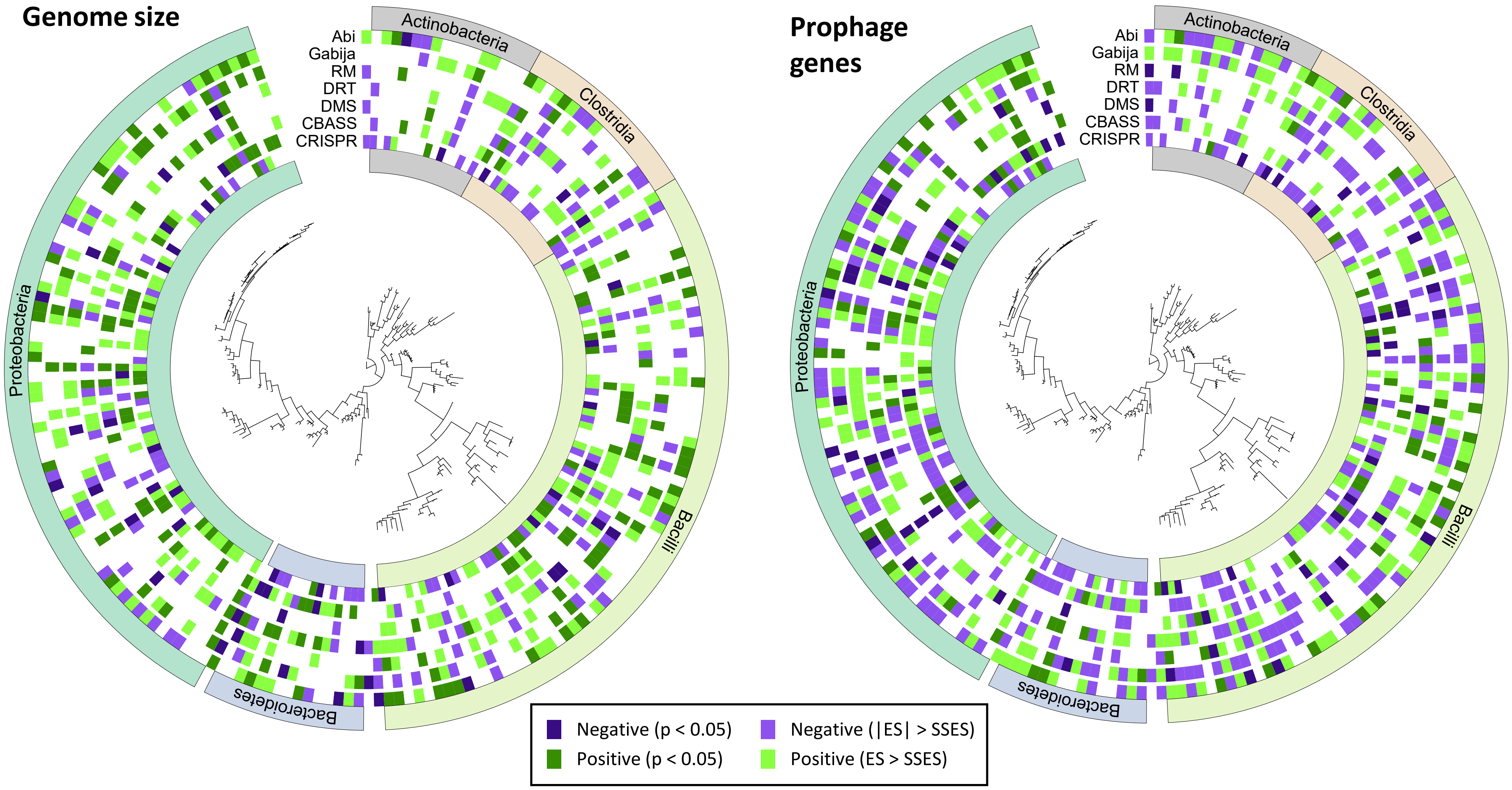


**Figure S4:** Taxonomic distribution of species displaying positive or negative associations between the presence of defense systems and the number of genes in the genome (left) or the number of marker genes for prophages (right).
