## Supplemental Figure S5 for "Timescale and genetic linkage explain the variable impact of defense systems on horizontal gene transfer"

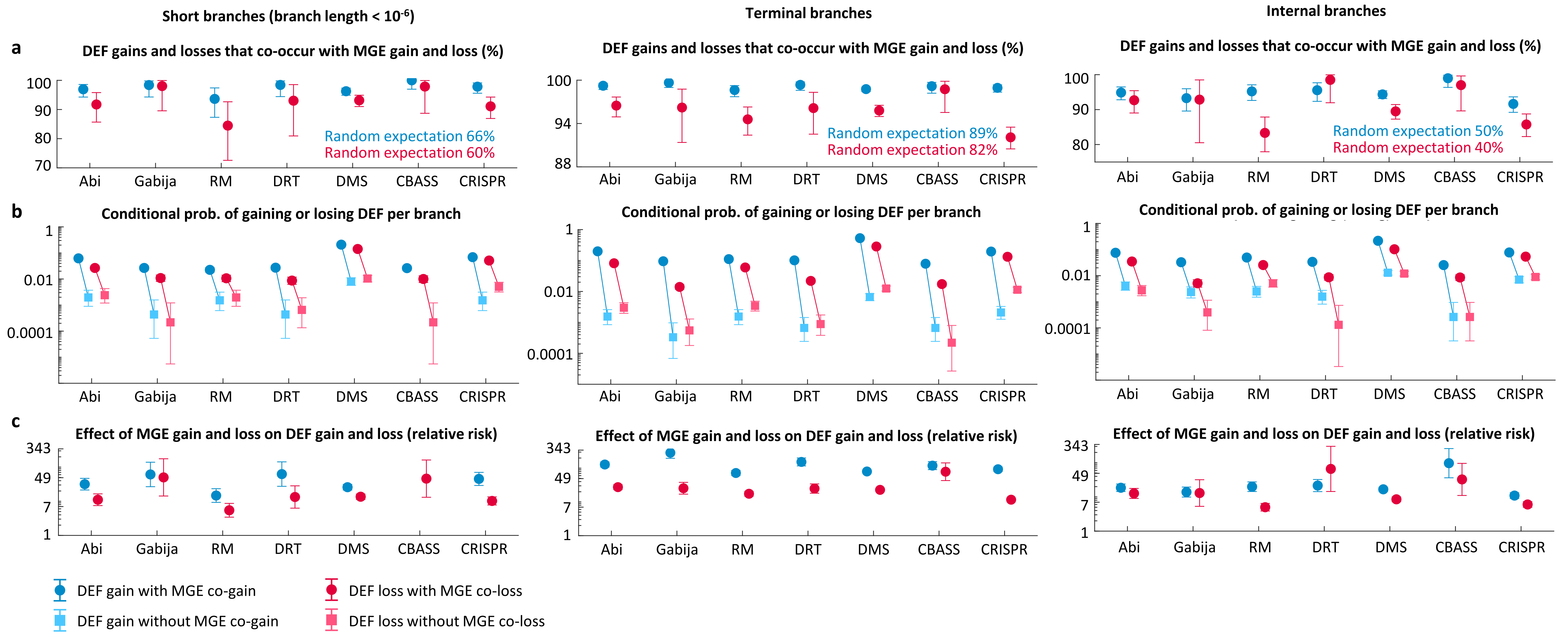


**Figure S5**: Co-occurrence of defense system gains and losses and MGE gains and losses in short branches (left), terminal branches (middle), and internal branches (right) of single-species phylogenetic trees. (a) Percentage of DEF gains (and losses) that occur in the same branch as an MGE gain (or loss). (b) Conditional probability of gaining (or losing) a defense system provided that an MGE is also gained (or lost) in the same branch, compared to the conditional probabilities when an MGE is not acquired (or lost) in the same branch. (c) Effect of MGE gain (or loss) on the per branch probability to acquire (or loss) a defense system, measured as a risk ratio. Error bars in (a) and (b) correspond to 95% confidence intervals based on the binomial distribution. Error bars in (c) indicate the 95% confidence intervals for the relative risk.
