## Supplemental Figure S6 for "Timescale and genetic linkage explain the variable impact of defense systems on horizontal gene transfer"

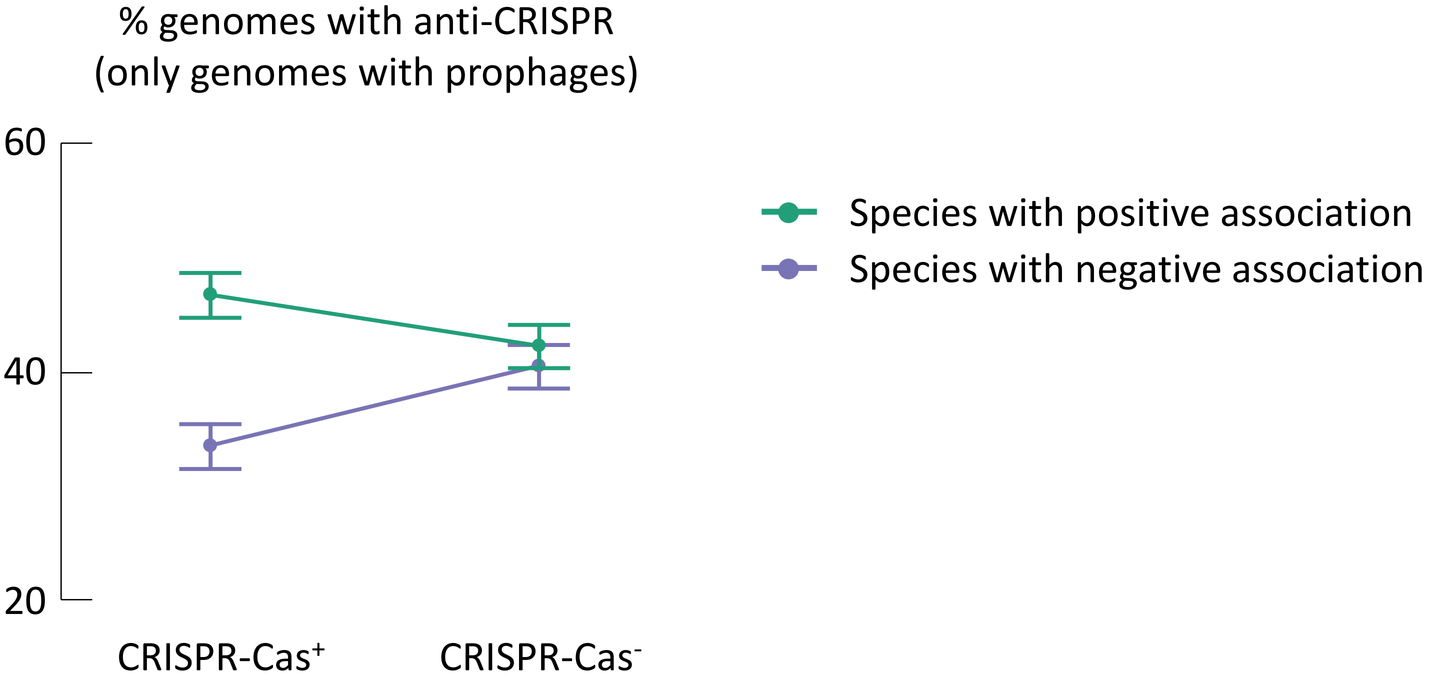


**Figure S6:** Prevalence of anti-CRISPR proteins (Acr) in genomes that do and do not contain CRISPR-Cas systems, separately calculated for species that display a positive or negative association between the presence of CRISPR-Cas and the number of genes from the mobilome (PGLMM, smallest significant effect size criterion). To control for the confounding effect of MGE prevalence on Acr, only genomes with prophages were considered. Whiskers represent 95% confidence intervals based on the binomial distribution.
